# COVID-19db linkage maps of cell surface proteins and transcription factors in immune cells

**DOI:** 10.1101/2022.12.14.520411

**Authors:** Koushul Ramjattun, Xiaojun Ma, Shou-Jiang Gao, Harinder Singh, Hatice Ulku Osmanbeyoglu

**Author notes:** These authors contributed equally: K.R., X.M.

## Abstract

The highly contagious SARS-CoV-2 and its associated disease (COVID-19) are a threat to global public health and economies. To develop effective treatments for COVID-19, we must understand the host cell types, cell states and regulators associated with infection and pathogenesis such as dysregulated transcription factors (TFs) and surface proteins, including signalling receptors. To link cell surface proteins with TFs, we recently developed SPaRTAN (Single-cell Proteomic and RNA-based Transcription factor Activity Network) by integrating parallel single-cell proteomic and transcriptomic data based on Cellular Indexing of Transcriptomes and Epitopes by sequencing (CITE-seq) and gene cis-regulatory information. We apply SPaRTAN to CITE-seq datasets from patients with varying degrees of COVID-19 severity and healthy controls to identify the associations between surface proteins and TFs in host immune cells. Here, we present COVID-19db of Immune Cell States (https://covid19db.streamlit.app/), a web server containing cell surface protein expression, SPaRTAN-inferred TF activities, and their associations with major host immune cell types. The data include four high-quality COVID-19 CITE-seq datasets with a toolset for userfriendly data analysis and visualization. We provide interactive surface protein and TF visualizations across major immune cell types for each dataset, allowing comparison between various patient severity groups for the discovery of potential therapeutic targets and diagnostic biomarkers.

## INTRODUCTION

SARS-CoV-2 has infected at least 638 million people worldwide resulting in over 6.62 million deaths as of November 22, 2022 [1]. There is clinical heterogeneity among infected individuals; ~15% are symptomatic, and <10% present with a severe form of the disease. However, 26.8% of hospitalized patients develop a critical disease. Despite the development of new COVID-19 therapies, they are still insufficient, especially with the rise of new variants of concern (VOCs). This is due largely to our incomplete understanding of the underlying mechanisms of the disease.

Both SARS-CoV and SARS-CoV-2 use the host ACE2 receptor protein enter the cell [2]. Other viral entry factors include FURIN, type II membrane serine protease (TMPRSS2), kidney injury molecule-1 (KIM-1), neuropilin-1 (NRP-1), cluster of differentiation 147 (CD147), dipeptidyl peptidase-4 (DPP4 or CD26), and C-type lectins (DC-SIGN or CD209, L-SIGN or CD209L, LSECtin or CLES4G, ASGR1, and CLEC10A). The viral infection sets off a cascade of interactions among host cellular proteins (e.g., signaling proteins and transcription factors). This complex network may restrict viral replication in host cells or, conversely, may be taken over by the virus for its replication.

SARS-CoV-2 requires a permissive cellular environment for its replication. It lacks its own transcription machinery and depends fully on host TFs to complete its life cycle. Further, SARS-CoV-2-infected host cells induce dramatic changes in the immune response. For example, T cells identify and eliminate infected cells, whereas B cells produce virus-specific antibodies. Hence, it is critical that we understand the diversity of host-specific T and B cells and whether this diversity affects vaccine efficacy. A better understanding of host immune cell states may also lead to the identification of accurate markers to predict the effects of infection on immune responses and identify novel therapeutic targets to treat COVID-19. In order to effectively treat the disease, we must elucidate the underlying immunological factors that cause critical COVID-19 illness and identify the associated immune cell states.

Single-cell-resolution genomics data allow us to characterize the full repertoire of SARS-CoV-2 infected cells and surrounding cells, which provide a high-resolution view of the states of the host immune cells. Cellular indexing of transcriptomes and epitopes by sequencing (CITE-seq) has been coupled with relatively sparse single-cell RNA sequencing (scRNA-seq) data and with robust detection of abundant and well-characterized surface proteins using barcoded antibodies [3, 4]. CITE-seq has the unique capability of reporting the transcriptomic and proteomic (surface protein) changes of various cell types in response to SARS-CoV-2 infection. We recently developed SPaRTAN (Single-cell Proteomic and RNA-based Transcription factor Activity Network) to mine the single-cell proteomic (scADT-seq) and corresponding scRNA-seq datasets obtained by CITE-seq. SPaRTAN links cell-specific expression of surface proteins with inferred transcription factor (TF) activities [5]. Although the cell surface phenotype of immune cells can be readily determined by flow cytometry, signaling pathways downstream of cell surface receptors/co-receptors drive changes in transcription and chromatin states. Therefore, to define cell states, it is important to connect the cell surface phenotype to the TF-mediated downstream transcriptional/transcriptomic phenotypes.

Several COVID-19-related CITE-seq datasets [6–9] have been generated, mostly from patient peripheral blood mononuclear cells (PBMCs). Connecting upstream signal transduction (e.g., cell surface receptors) to downstream TFs at a single-cell level could identify cell states and characterize SARS-CoV-2-infected cells and surrounding cells to determine how they contribute to COVID-19 pathogenesis. We applied SPaRTAN to COVID-19 CITE-seq datasets to predict the coupling of surface receptors with TFs across host immune cells (**Figure 1, Table 1**). To support the research community in investigating and using our analyses, we created the web resource for COVID-19db linkage maps of cell surface proteins and transcription factors in immune cells. For each study, we provide 1) surface protein expression, 2) SPaRTAN-predicted TF activities for each cell type per patient for each dataset, 3) COVID-19-related publications, and 4) protein-protein interaction (PPI) networks, pathways, and drugs associated with surface protein or TFs of interest. Our web resource provides unique and valuable data for testing hypotheses and identifying surface proteins associated TFs that could facilitate the understanding of COVID-19 pathogenesis and identify novel therapies.

**Figure 1:**
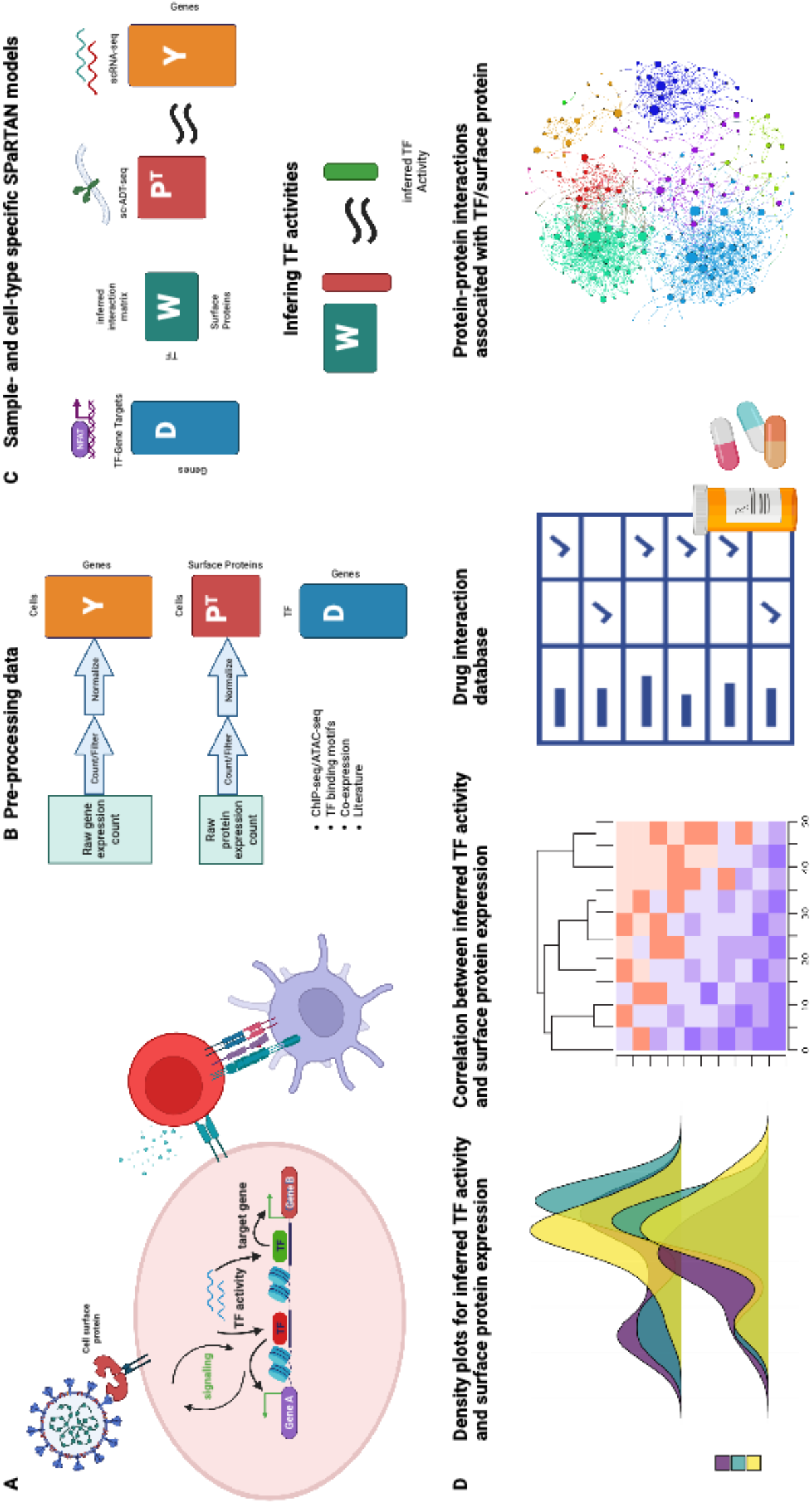
An integrative computational model linking cell surface receptors to TFs. **(A)** Our integrative model (SPaRTAN, Single-cell Proteomic and RNA-based Transcription factor Activity Network) infers the flow of information from cell surface receptors to transcription factors (TFs) to target genes by learning the interactions between cell surface receptors and TFs that best predict target gene expression. The general SPaRTAN workflow starting from **(B)** data preprocessing, **(C)** building sample-specific and cell type-specific SPaRTAN models and inferring TF activities based on the model, and **(D)** downstream analysis (e.g., TF/surface protein density/ridge plots across disease subgroups, correlation heatmaps between TF activity and surface protein expression per sample, relevant drug targets, and PPI networks). Created with BioRender.com.

**Table 1.**
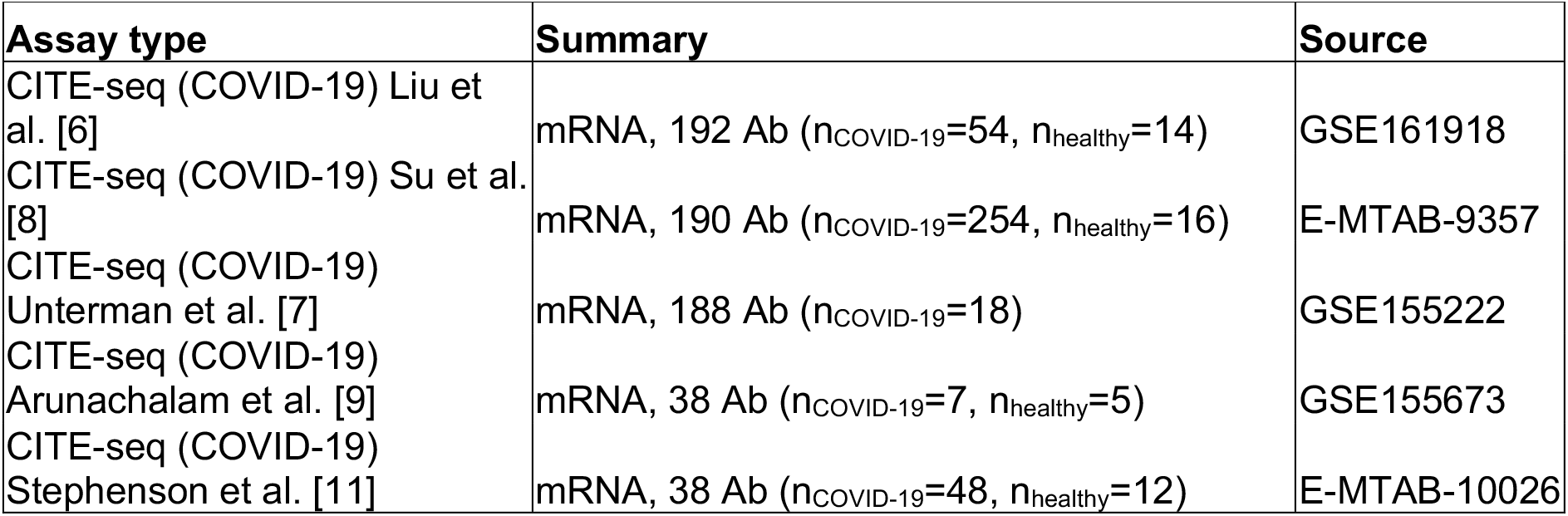
Five high-αualitv COVID-19 CITE-seq datasets included in our webserver

| Assay type | Summary | Source |
| --- | --- | --- |
| CITE-seq (COVID-19) Liu et al. [6] | mRNA, 192 Ab ($n_{\text{COVID-19}}=54$, $n_{\text{healthy}}=14$) | GSE161918 |
| CITE-seq (COVID-19) Su et al. [8] | mRNA, 190 Ab ($n_{\text{COVID-19}}=254$, $n_{\text{healthy}}=16$) | E-MTAB-9357 |
| CITE-seq (COVID-19) Unterman et al. [7] | mRNA, 188 Ab ($n_{\text{COVID-19}}=18$) | GSE155222 |
| CITE-seq (COVID-19) Arunachalam et al. [9] | mRNA, 38 Ab ($n_{\text{COVID-19}}=7$, $n_{\text{healthy}}=5$) | GSE155673 |
| CITE-seq (COVID-19) Stephenson et al. [11] | mRNA, 38 Ab ($n_{\text{COVID-19}}=48$, $n_{\text{healthy}}=12$) | E-MTAB-10026 |

## MATERIAL AND METHODS

### Data resources for building SPaRTAN models

We downloaded COVID-19 CITE-seq datasets from the website at https://atlas.fredhutch.org/fredhutch/covid as well as from original publications [6–11] (**Table 1**). We collected the surface protein expression matrix of the raw count (scADT-seq), gene expression matrix of the raw count (scRNA-seq), and sample information, including the patient ID, disease (COVID-19) subgroup, and cell type annotation for each dataset from original studies. The surface proteins included in the database are identified by their UniProt names or HUGO Gene Nomenclature Committee gene names. To stratify patients for analysis, we focused on disease subgroups that described each patient’s severity level by symptoms. Disease severity groups, based on the National Institutes of Health guidelines **[12],** included healthy donors, COVID-19 (mild), COVID-19 (moderate), COVID-19 (severe), and COVID-19 (critical) patients. For the GSE155224 dataset, we included COVID-19 (stable) and COVID-19 (progressive), as indicated in the original study [7].

We created the TF-target gene prior matrix for SPaRTAN analysis to define a candidate set of associations between TFs and target genes. To determine the set of TFs that potentially regulates each gene, we downloaded curated TF-target gene interactions from hTFtarget [13]. hTFtarget includes curated TF-target genes from ChIP-Seq data (7,190 experiments related to 659 TFs) across 569 conditions (399 types of cell lines, 129 tissue or cell types, and 141 treatments).

### Data preprocessing for SPaRTAN analysis

Normalization and initial explanatory analysis of CITE-seq datasets were performed using the Seurat R package version 4.0.4 [14]. We excluded cells with less than 300 expressed genes and 1,000 detected molecules to ensure cell quality. We also excluded cells with more than 5,000 expressed genes to avoid doublets. Antibody-derived tags (ADTs) were normalized using a centered log ratio (CLR) transformation across cells, and scRNA-seq data were log-normalized using a size factor of 10,000 molecules for each cell.

For training sample-specific and cell type-specific SPaRTAN models, we sorted each dataset by sample id and further grouped cells into six major immune cell types (monocytes, B, natural killer [NK], CD4^+^ T, CD8^+^ T, and dendritic cells). For each sample and cell type, we excluded genes that were expressed (>0) in less than 3% of cells; and surface proteins that were expressed (>0) in less than 15% of cells.

We also excluded TFs in the TF-target gene interaction matrix that were not expressed in all cells for a particular cell type for each patient. We filtered this TF-target gene interaction matrix so that each gene has at least five TFs, and each TF has at least 10 target genes but no more than 80% of all genes. The source code for processing each CITE-seq dataset and constructing the TF-target gene prior matrix is deposited at the Github repository https://github.com/osmanbeyoglulab/covid19_webapp.

### Training sample-specific and cell type-specific SPaRTAN models

SPaRTAN uses the expression of surface proteins (based on scADT-seq) as a proxy for their activities (**Figure 1B)**. Signaling via these proteins converges on TFs that, in turn, regulate the expression of their target genes (based on scRNA-seq, TF-target gene priors). Formally, we use a regularized bilinear regression algorithm called affinity regression (AR) [15–17], which provides a statistical framework when the observed data can be explained as interactions between two kinds of inputs. Here, SPaRTAN establishes an interaction matrix (**W**) between surface proteins (**P**) and TFs (**D**) that predicts target gene expression (**Y**) (**Figure 1C**). The model captures statistical relationships between surface proteins, TFs, and gene expression. We used the trained interaction matrix (**W**) to predict TF activity from the surface protein expression profile of a cell. We trained sample-specific and cell type-specific SPaRTAN models [18] and predicted cell-specific TF activities for each CITE-seq dataset [6–10]. We also calculated the correlation between surface protein expression and TF activities across cells. Inferred TF activities and surface protein expression can provide insights into the states of host immune cells.

### TF-target gene analysis

To identify the associations between SPaRTAN-inferred TF activities and target gene expression for a cell type of interest, we computed Spearman correlation coefficients (rho) between (inferred) TF activity and gene expression for each TF-gene pair per sample. For clarity, pairwise absolute Spearman correlation values below 0.3 in all samples were excluded. For each TF of interest, we also downloaded a list of target genes using a custom hTFtarget API and collected the result along with the epigenomic states into a comprehensive table.

### Linking surface proteins with TFs

To identify the associations between inferred TF activities and surface protein expression at a single-cell level, we computed Spearman correlation coefficients (rho) between (inferred) TF activity and surface protein expression for each TF-surface protein pair within each cell type per sample. We used a heatmap to display correlations. For clarity, pairwise Spearman correlation values below 0.6 in all samples were excluded.

### TFs and surface proteins reported in COVID-19-related literature

We downloaded the LitCovid [19] dataset to search for the latest COVID-19 literature relevant to the associated TFs and surface proteins in our study. Briefly, the LitCovid dataset comprises PubMed-indexed articles related to SARS-CoV-2 [19]. The dataset currently contains 302,986 articles that grow daily, making it a comprehensive, up-to-date resource for COVID-19 researchers.

### PPIs associated with TFs and surface proteins

To provide a visual model of high-confidence genes that interact with TFs and surface proteins of interest, we utilized the protein-protein interaction (PPI) network derived from the STRING database (StringDB) [20] (confidence score > 400) that contains protein-PPIs from experimental and computational sources. Using the StringDB native application programming interface (API), we queried the database for interactions involving selected TFs or surface proteins using the Python requests library. The number of interactors can be manually set by the user on the PPI tab. The width of the edges corresponds to StringDB scores. Only connected nodes are plotted.

As a supplementary resource, we also included predicted interactions from Wiki-Corona [21]. Wiki-Corona is a resource centered on human PPIs that may be involved in SARS-CoV-2 infection. Using a custom-built API wrapper, we also queried the Wiki-Corona website to retrieve information about pathways associated with proteins/TFs of interest, any associated diseases, and drugs that bind to relevant factors.

### Mapping drugs to target TFs and surface proteins

The drugs and their annotations were imported from the Drug Repurposing Hub [22] from the ConnectivityMap (CMap) database [23].

### Statistical analysis and visualization

The distributions of the expression of surface proteins and TF activity ranks were organized by disease subgroups. Specifically, we used ridge density plots for each dataset. The significance of the difference between the disease subgroups in each cell type was evaluated through the Mann-Whitney U test. Graphs were generated using ggplot2 (version: 3.3.5), pheatmap (version: 1.0.12), and ggridges (version: 0.5.3) packages. For general data analysis and manipulation, Seurat (version: 4.0.4), SeuratDisk (version: 0.0.0.9020), rstatix (version: 0.7.0), dplyr (version: 2.1.1), data.table (version: 1.14.2) and tidyr (version: 1.2.0) were used.

### Web portal for the database

We built the COVID-19db of Immune Cell States web portal to present user-friendly analysis results. All the processed and annotated datasets can be searched, visualized, and downloaded from the web portal **(Figure 2**). The back end of the portal was implemented in Python, while the front end was written in TypeScript (version: 4.9.3) via the Streamlit (version: 1.14.0) open-source framework. All the charts were generated by in-house Python (version: 3.9.0) and R scripts. COVID-19db of Immune Cell States database is deployed with the Streamlit share server and is freely available (http://covid19db.streamlitapp.com) without registration or login. All the functions of the database have been tested in Google Chrome and Apple Safari browsers. Static figures and logos were made using BioRender. (www.biorender.com)

**Figure 2:**
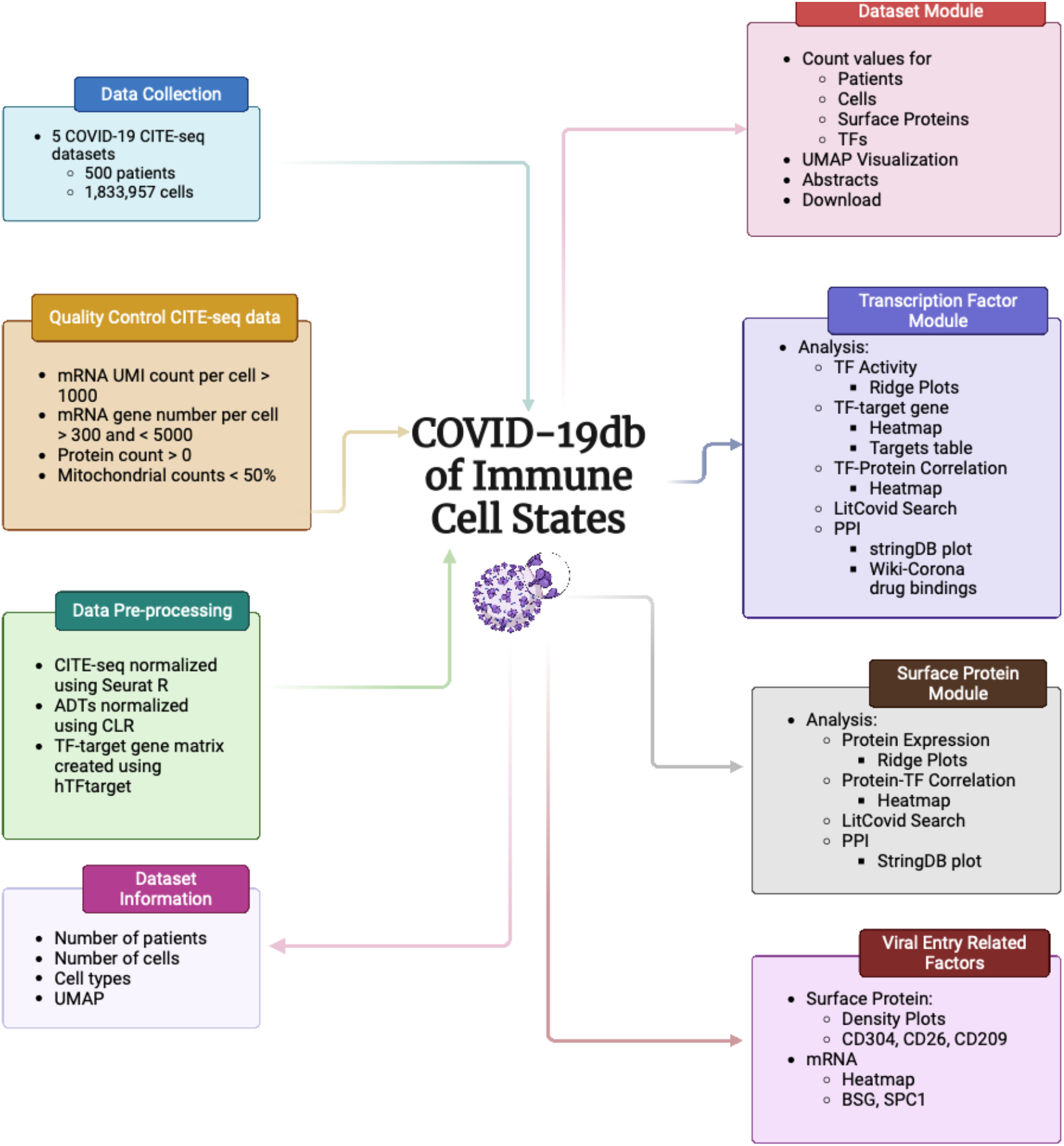
Overview of the COVID-19db linkage maps of cell surface proteins and transcription factors in immune cells workflow and features. We curated COVID-19 CITE-seq datasets from GEO or Array Express databases. All datasets were then uniformly processed with a standardized workflow, including quality control. Cell type annotations were retrieved from original studies. Each dataset is displayed with relevant study information, including the number of patients and cells, patient severity groups, as well as the number of surface proteins and TFs associated with the related study. In the Dataset module, we provide three modules, surface protein exploration, TF exploration, and viral entry-related factors exploration. In the surface protein module, we provide visualization of a single surface protein for a chosen dataset and a specific cell type across patient severity groups, as well as correlation with TFs. In the TF module, we provide visualizations of cell typespecific TF activity across disease subgroups for chosen datasets as well as a correlated surface proteins. In the viral entry-related factors module, we provide visualization of viral entry related factor protein expression or gene expression depending on the availability in the selected dataset of the chosen datasets for a cell type across patient severity groups as well as correlated TFs.

## RESULTS

We applied SPaRTAN to publicly available PBMC CITE-seq datasets of healthy individuals and patients with various severities of COVID-19 to identify the dysregulated critical TFs downstream of surface receptors that drive the various cell states for each major immune cell type. Then, we created a web resource that provides a user-friendly interface for systematically visualizing, searching, and downloading data on surface protein expression, TF activity, and their associations with various immune cell types. Accessing these data allows for a fast, flexible, and comprehensive exploration of host immune cell types that are altered in COVID-9 patients. The current database contains five CITE-seq datasets for six major immune cell types, including monocytes, B, NK, CD4^+^ T, CD8^+^ T, and dendritic cells (DC). Most datasets contain samples from healthy donors to provide baseline expression levels for immune cells (**Table 1**). The overall curation workflow of web resource is illustrated in **Figure 2**.

### Dataset browser

In the dataset browser module, users can access information about the various COVID-19 CITE-seq datasets available for further analysis by subsequent modules. For ease of presentation, the projects are referred to by their first-listed author, GEO/Array Express accession number, and PubMed Unique Identifier (PMID). This dataset browser module provides quick, convenient information on the number of patients and the total number of cells per dataset, as well as the number of surface proteins and TFs used in the SPaRTAN analysis. For each selected dataset, additional information, such as the abstract of the original study, data download links, and the uniform manifold approximation and projection (UMAP) plot for visualization, is provided.

### Analysis by surface proteins

Each dataset provides the user with the available immune cell types and surface proteins. The analysis by surface proteins module allows users to explore expression distributions for a surface protein of interest and determine correlations between surface protein expression and SPaRTAN-inferred TF activities for each sample by immune cell type. The distribution of the selected protein expression in the selected cell type is computed dynamically and shown for the various disease subgroups as a ridge plot. For a selected surface protein, a COVID-19-relevant literature search based on the LitCovid database [19], PPIs based on StringDB [20], and relevant pathways and drugs based on Wiki-Corona [21] are displayed (e.g., **Figure 3**).

**Figure 3:**
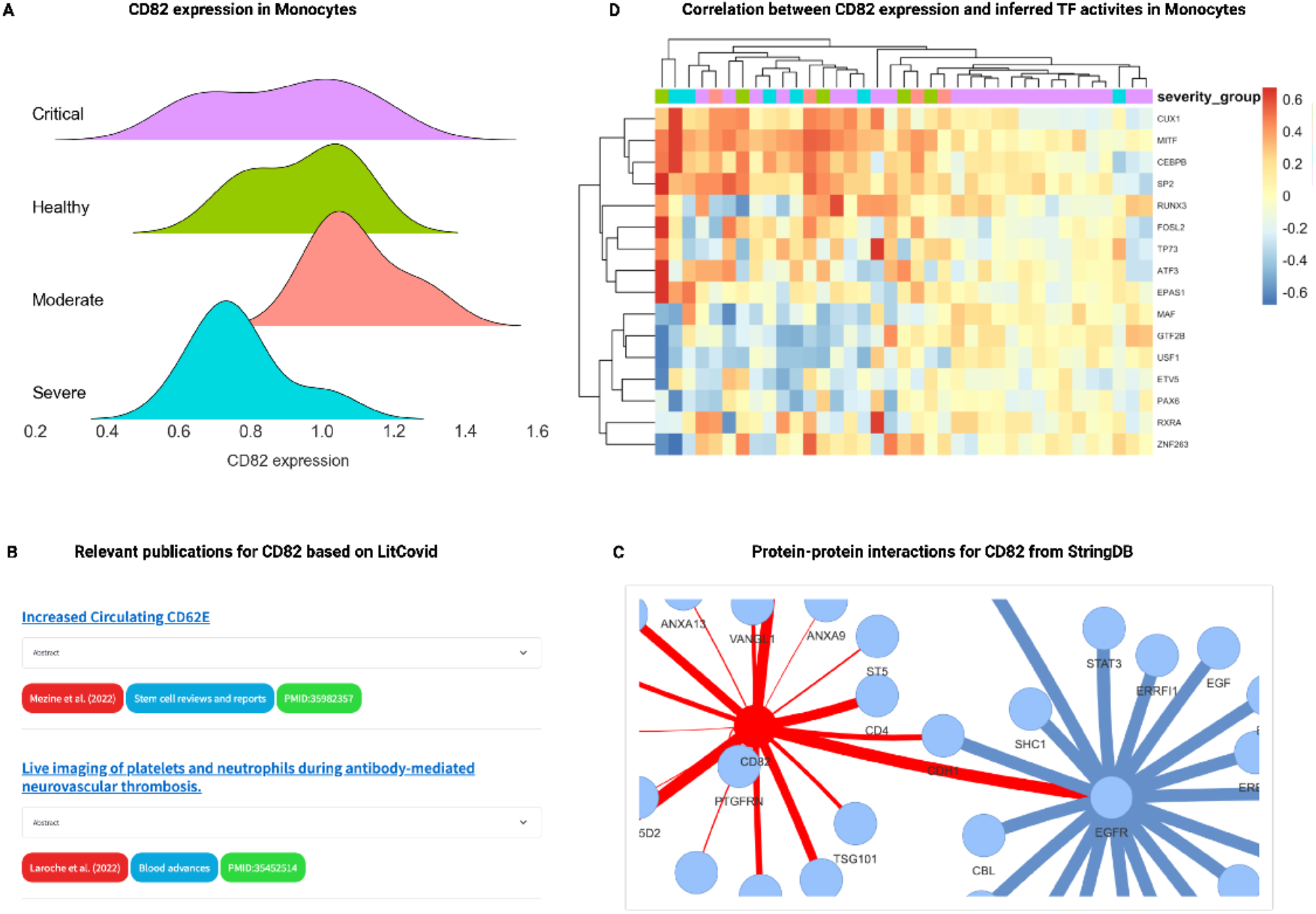
Analysis by a surface protein. **(A)** A user-specified surface protein is dynamically displayed in grouped ridge plots by disease subgroup. **(B)** COVID-19 publications related to the surface protein of interest are generated by the LitCovid database. **(C)** PPI networks associated with the selected surface protein are generated by the StringDB, and pathways and drugs are generated by Wiki-Corona. **(D)** A heatmap revealing correlations between surface protein expression and inferred TF activities of cells (rows) for samples in a selected cell type. For clarity, TFs with pairwise Spearman correlation values below 0.3 in at least one sample are excluded.

### Analysis by transcription factors

For each selected cell type-TF pair, the analysis by transcription factor module allows users to explore TF activity distribution based on its rank by disease subgroup, which is shown as a ridge plot. Users can visualize correlations between the activity of the TF of interest and surface protein expression and target gene expression across samples for a selected cell type. In addition, the module provides a COVID-19-relevant literature search based on the LitCovid database [19], PPIs based on StringDB [20], and relevant pathways and drugs based on Wiki-Corona [21] (e.g., **Figure 4**).

**Figure 4.**
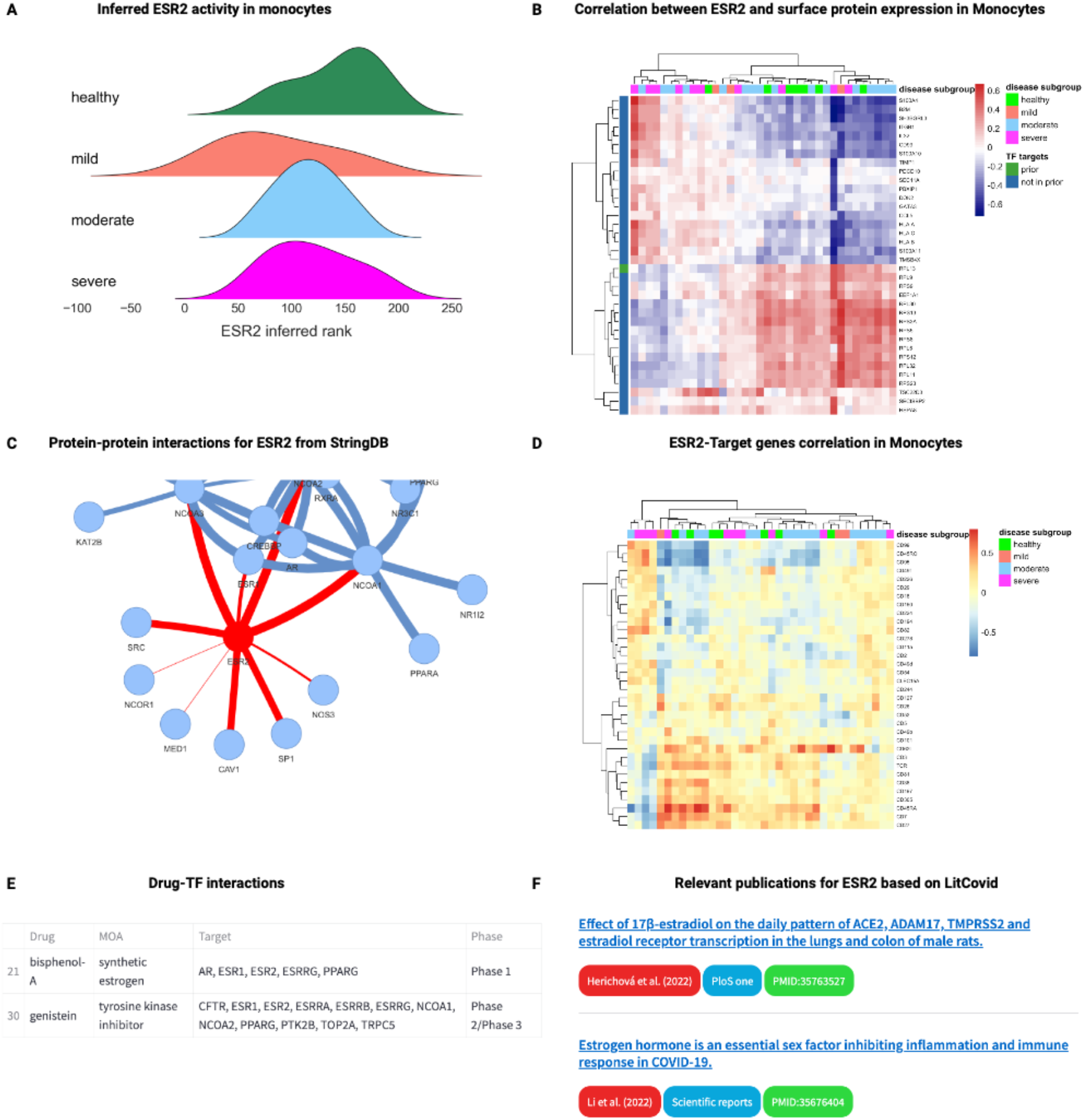
Analysis by a TF. **(A)** Predicted TF activity rank based on a SPaRTAN analysis specified by the user can be dynamically displayed in grouped ridge plots by disease subgroup. **(B)** A heatmap revealing correlations between inferred TF activity and surface protein expression across cells (rows) for samples in a selected cell type. For clarity, surface proteins with pairwise Spearman correlation values with TFs below 0.3 in at least one sample are excluded. **(C)** PPI network associated with selected surface protein generated by the StringDB, and pathways and drugs are generated by Wiki-Corona. **(D)** Heatmap based on the subset of potential context-specific target genes in selected cell type where gene expression correlated with SPaRTAN-inferred TF activity across patient groups (target genes with |*ρ*| > 0.3 shown). Green labels indicate that the target gene was in prior TF-target gene interactions based on the hTFtarget dataset, and blue labels indicate the target gene was not in prior TF-target gene interactions. **(E)** Available TF-drug interactions **(F)** COVID-19 publications related to the surface protein of interest are generated by the LitCovid database.

### Analysis by viral entry-related factors

For each dataset, users are provided with genes relevant to viral entry, including *NRP1/CD304, DPP4/CD26, DC-SIGN/CD209, CD147/BSG*, or *FURIN/SPC*. Users can explore the protein or gene expression distribution patterns for these genes for a particular cell type, depending on availability. In addition, the module provides a COVID-19-relevant literature search based on the LitCovid database [19], PPIs based on StringDB [20], and relevant pathways and drugs based on Wiki-Corona [21].

## DISCUSSION

The COVID-19db linkage maps of cell surface proteins and transcription factors in immune cells web portal facilitates the exploration of the cell state-specific regulators, including surface proteins and TFs. These can be used by experimental scientists to determine commonalities and differences in cell context-specific TF and surface receptors in healthy individuals versus COVID-19 patients with varying levels of severity of the disease.

In this study, we applied SPaRTAN to COVID-19 CITE-seq datasets, linking surface proteins to TFs for each host immune cell type. Using datasets from four studies, we generated a public database, COVID-19db linkage maps of cell surface proteins and transcription factors in immune cells, which provides analyses of surface proteins and TFs across immune cell types for COVID-19 patients and healthy donors. Additionally, we provide an example of a set of analyses that we performed using our web source.

Despite an explosive increase in the generation of COVID-19-related data, knowledge of the mechanistic basis of the host cellular response to SARS-CoV-2 infection is limited. Singlecell characterization of cell types will improve our understanding of virus-host interactions and the susceptibility of COVID-19 patients with chronic respiratory disease to severe disease. This information can provide mechanistic insights concerning the immune response to SARS-CoV-2, identify potential biomarkers, and yield therapeutic targets that specifically benefit COVID-19 patients with pre-existing respiratory conditions.

Limitations: It is difficult to compare the results from different studies because they usually have different sets of surface proteins. Also, because blood is easy to collect from patients, the tissue in most COVID-19 CITE-seq studies is limited to PBMCs. When CITE-seq data from tissues other than blood are available, we will update the website with these data for a better understanding of the multi-organ impact of COVID-19. Moreover, linkages that SPARTAN infers can imply one way or reciprocal crosstalk between a cell surface protein and a TF (e.g. (i) TF induces or represses cell surface protein (ii) Cell surface protein signalling alters TF activity and (iii) TF and cell surface protein mutually reinforce each other’s activity/expression).

In summary, COVID-19db linkage maps of cell surface proteins and transcription factors in immune cells is a useful repository for COVID-19 researchers. It provides a user-friendly web resource for interactive TF and surface protein visualizations for each immune cell type. Our webpage will allow immunologists to study gene regulation and immune signaling. In the future, we will continue to improve our webpage by maintaining current web resources and regularly adding and integrating new datasets. As more public COVID-19 CITE-seq data are available, we anticipate that this continued development and maintenance of the web resource will benefit the broader research community.

## CODE AVAILABILITY

The code for our web resource is available from http://github.com/osmanbeyoglulab/covid19_webapp

## ACKNOWLEDGEMENT

The authors acknowledge Sanghoon Lee and April Sagan for the helpful discussion and suggestions on the website. The authors acknowledge the authors from published studies to share their data on COVID-19 CITE-seq profiling. Data analyses in this research were supported by the University of Pittsburgh Center for Research Computing and the Extreme Science and Engineering Discovery Environment (XSEDE), which is supported by National Science Foundation grant number OCI-1053575. Specifically, it used the Bridges2 system, which is supported by NSF award number ACI-1445606, at the Pittsburgh Supercomputing Center.

